# A decade of empirical research on research integrity: what have we (not) looked at?

**DOI:** 10.1101/567263

**Authors:** Noémie Aubert Bonn, Wim Pinxten

## Abstract

In the past decades, increasing visibility of research misconduct scandals created momentum for discourses on research integrity to such an extent that the topic became a field of research itself. Yet, a comprehensive overview of research in the field is still missing. Here we describe methods, trends, publishing patterns, and impact of a decade of research on research integrity.

To give a comprehensive overview of research on research integrity, we first systematically searched SCOPUS, Web of Science, and PubMed for relevant articles published in English between 2005 and 2015. We then classified each relevant article according to its topic, several methodological characteristics, its general focus and findings, and its citation impact.

We included 986 articles in our analysis. We found that the body of literature on research integrity is growing in importance, and that the field is still largely dominated by non-empirical publications. Within the bulk of empirical records (N=342), researchers and students are most often studied, but other actors and the social context in which they interact, seem to be overlooked. The few empirical articles that examined determinants of misconduct found that problems from the research system (e.g., pressure, competition) were most likely to cause inadequate research practices. Paradoxically, the majority of empirical articles proposing approaches to foster integrity focused on techniques to build researchers’ awareness and compliance rather than techniques to change the research system.

Our review highlights the areas, methods, and actors favoured in research on research integrity, and reveals a few blindspots. Involving non-researchers and reconnecting what is known to the approaches investigated may be the first step to generate executable knowledge that will allow us to increase the success of future approaches.

**A word from the authors:** We find important to mention that this manuscript underwent peer review and was rejected from the following journals:

- PLOS ONE
  - Submitted 19^th^ December 2017
  - Peer-review and rejection received 26^th^ June 2018.
- Journal of Science and Engineering Ethics (JSEE)
  - Submitted 18^th^ August 2018
  - Peer-review response with major revision request received 9^th^ September 2019
  - Revision submitted 24^th^ October 218
  - Rejection received 27^th^ December 2018

We regret not having submitted this preprint before our first submission. Nonetheless, now after over one year in submission processes, we thought that we should make this manuscript and its data available as a pre-print before undergoing further submissions.

In order to promote transparency however, we asked both journal whether anonymous reviews could be added alongside this pre-print to ensure that readers are informed of the issues that disqualified our manuscript.

PLOS ONE agreed for us to share the anonymous reviews which are now available — together with our itemized changes and responses — in the ‘Online Resource 5 – Peer Review Report’. We thank the editors of the Journal of Science and Engineering Ethics and the integrity team of Springer Nature for thoroughly discussing our request, but unfortunately, given the closed peer review policy at Springer Nature, we were unable to provide information about the peer review from the Journal of Science and Engineering Ethics.

We advise our readers to look at the peer-review and be aware of the challenges and limitations attached with our work. Of course, we welcome comments and contributions to make our work better.

Sincerely,

Noémie Aubert Bonn and Wim Pinxten

## Introduction

Research integrity (RI) has been part of the scientific discourse for many years, and evolved to a topic of research itself over the past 20 years. Research on RI highlighted that research misconduct comes in many forms (De Vries et al. 2006), occurs more often than was initially thought, and that questionable research practices (QRP)—also referred to as *detrimental research practices*, practices outside the realm of misconduct which still risk damaging the scientific output—are far from rare (Fanelli 2009; Pupovac and Fanelli 2014).

In 1999, one of the first paper setting the agenda for research on RI concluded with the following words:

> *Over the last decade, researchers and research institution have made significant strides toward restoring [trust in science] by actively confronting misconduct. […] With so much accomplished, the time is right to see whether the policies we have put in place, the funds and time spent, have made a difference. Have we achieved levels of integrity in research that are acceptable?* (Steneck 1999, p. 173)

Now nearly two decades later, this call for research on RI seems to have been heard. Scientific literature on research integrity and research misconduct increased exponentially, broad scale funding and consortiums have been established to enable more research on the topic (e.g., the European Commission Horizon 2020 contributed an impressive 20 million euros in projects on RI since 2015), attendance to the last *World Conference on Research Integrity* exceeded 900, and some institutions are starting build departments with PhD students specializing on the topic.

Notwithstanding this growing interest for research on RI and misconduct, it is unclear how the potential to identify and quantify the problems, to highlight and understand determinants of bad science, and to assess and propose approaches that foster integrity and prevent misconduct have been employed. To provide better insight in the field, we analysed published research on RI. The goal of this analysis was twofold. On the one hand, we aimed to understand how researchers focusing on RI perform research (i.e.: which methods are used, which stakeholders are studied, and which topics are most investigated). On the other hand, we aimed to document gaps of knowledge to inform future research endeavours.

## Methods

Studying research on RI is methodologically challenging. Researchers from many different fields address the topic in different ways. There is poor consistency in how the scope of RI is delimited (e.g., Is research ethics part of integrity? Is academic integrity only targeting students?) and in the choice of journals or article formats. For example, the empirical piece of Brian Martinson and colleagues (2005)—widely recognised as a cornerstone in research on RI—was published as a ‘Commentary’ in Nature, is currently being classified as a ‘Note’ in Scopus and as ‘Editorial material’ in the Web of Science. Consequently, systematic searches for relevant empirical works on research integrity have serious blind spots if the sample is kept manageable.

We are aware that, despite all efforts to gather a manageable sample of the highest possible relevance, the choices we made towards our search strategy unavoidably come at a cost (e.g., not including the Martinson et al. paper, and unavoidably several others). We ensured that such costs are transparently reflected throughout this paper.

To characterize the broad spectrum of research on research integrity, we performed an analysis of the literature on RI published in English between 2005 and 2015. Our analysis differs from a typical literature review: we classified several variables beyond the findings of the included articles (e.g., publication year, impact metrics, geographical distribution, and several methodology characteristics) and analyzed the relationship between such variables. Consequently, our findings do not describe what is known about specific aspects of RI, but rather provide an overview of how research on RI is performed and published in order to highlight the areas or actors where/with whom most research is done and areas or actors current research might have overlooked.

We used three major bibliographic databases to find relevant literature on RI: SCOPUS, Web of Knowledge, and PubMed. We performed and adapted our search between February and April 2017 for SCOPUS, between October and November 2017 for Web of Science and PubMed, and in February 2019 to add the terms ‘scientific fraud’ and ‘research fraud’ as recommended by reviewers. We extracted all results in an Excel sheet, which is available as tab delimited in ‘Online Resource 3 - General data’. We only kept the records present on the sheet for further analyses. The complete study flow diagram with inclusion/exclusion counts and search queries may be seen in Fig 1.

**Figure 1.**
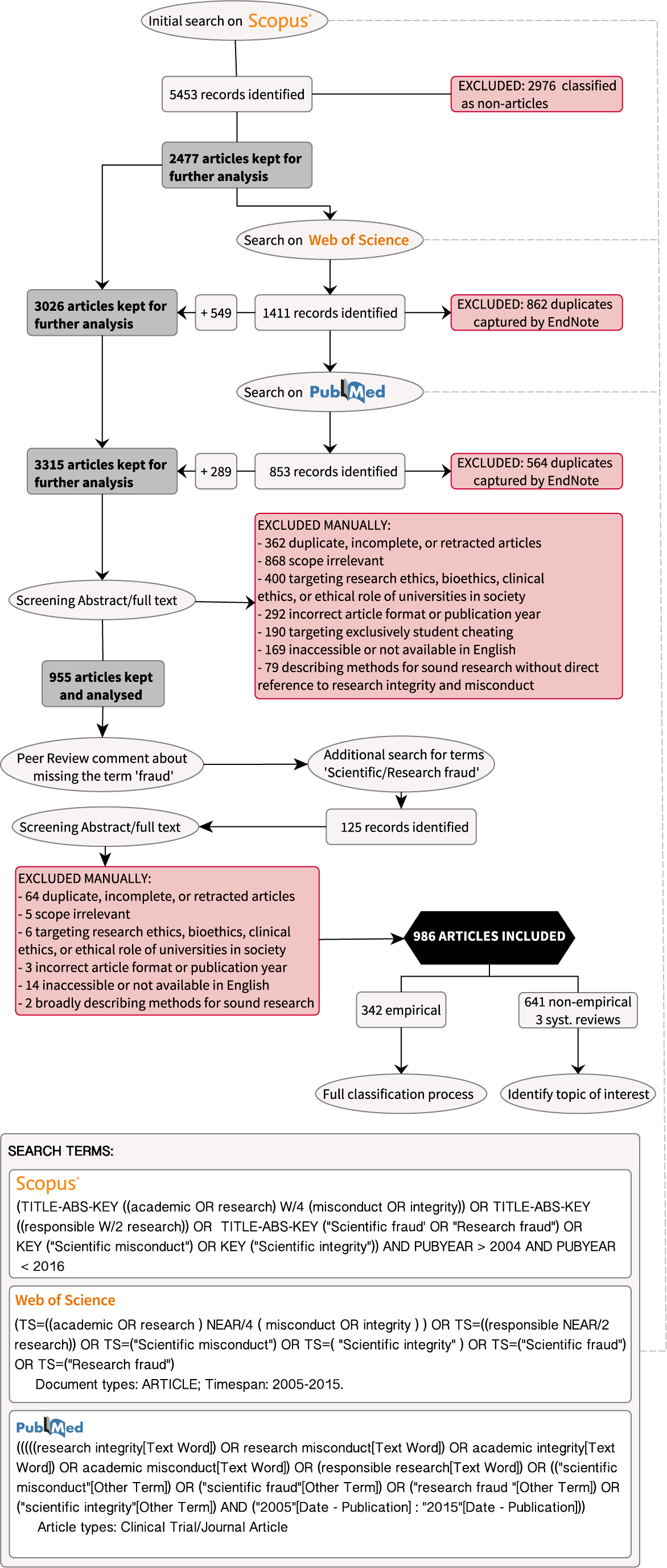
Study flow diagram.

In summary, our queries screened titles, abstracts, and keywords of published literature for any mention of ‘academic misconduct’, ‘academic integrity’, ‘research misconduct’, ‘research integrity’ (or any expression of six words or less containing such terms), ‘responsible research’ (or any expression of four words or less containing these two terms), ‘scientific integrity’, ‘scientific misconduct’, ‘scientific fraud’, and ‘research fraud’. We chose these keywords after a few adaptations as we believed that they would provide a broad and yet specific enough overview of works that have been published on RI. Having worked in the field of RI in non-English speaking countries for some time, we purposively included the expression ‘academic misconduct’ despite its more direct relationship to student cheating to allow capturing articles which might have used the term differently to refer to research misconduct (Aubert Bonn et al., 2018).

We did not include papers relating to the ethical care of animals in research. Beyond papers whose scope was directly irrelevant, we also excluded several themes which were related, but not directly linked to RI, namely (i) academic integrity or cheating limited to undergraduate students, or with no apparent extension to RI in the discussion and the abstract of the paper (later referred as *cheating (exclusively)*); (ii) research ethics looking at the protection of human participants; (iii) clinical ethics or bioethics; (iv) responsible research innovations focusing on societal concerns of research discovery (ii-iv later referred as *Research ethics, bioethics, clinical ethics, or ethical role of universities in society*); and (v) techniques meant to improve the validity of research, but devoid of direct reference to QRP, misconduct, or integrity (later referred as *methods and tools*).

### Classification process

To build the classifications for our research, we used an inductive process based on the findings from the first set of papers retrieved (i.e., the SCOPUS search). An inductive process means that we started with the general goal of *describing research*, and that we decided on which categories and classification options we should include based on what we found in the abstracts and papers assessed. For this analysis, NAB built the search, retrieved the literature, selected articles to be included, and inductively classified the articles in categories. WP helped refine and simplify the categories, revised individual papers which were ambiguous, and provided assistance on the specific wording used for the categories.

A full description of the inductive process that led to our final categories is available in ‘Online Resource 1 – Building the classifications’. The final categories and classification options and their definitions are listed in Table 1. The full description of each category is included in ‘Online Resource 2 – Instructions for Use’.

**Table 1.** Classification categories found inductively and used in our analyses. A more detailed description of the classification process and how we developed the categories inductively is available in the ‘Online Resource 1 – Building the classifications’. For each relevant paper, we first noted the topic of interest. We then identified the paper as either empirical or not empirical. For empirical papers, we further identified the general methodology used, the studied population, the source where the data came from, the focus on research integrity issues, and the general research objective.

| TOPIC OF INTEREST | TYPE OF RESEARCH | GENERAL METHODOLOGY | STUDIED POPULATION | SOURCE OF DATA | FOCUS | OBJECTIVE |
| --- | --- | --- | --- | --- | --- | --- |
| • QRP and misconduct (general)<br>• Research integrity (general)<br>• RCR training, education and mentorship<br>• Cheating and academic misconduct<br>• Publication ethics<br>• Authorship and collaborations<br>• Plagiarism<br>• Conflicts of interest<br>• Guidelines and policies<br>• Peer Review<br>• Research infrastructures and environments<br>• REC/IRB<br>• Allegations, sanctions, disclosure of cases<br>• Whistleblowing<br>• CV and application misrepresentation<br>• Research on research integrity | • Empirical | • Surveys, Interviews or focus groups<br>• Content and textual analysis<br>• Bibliometric study<br>• Investigation or forensic analysis<br>• Tool building or validation<br>• Combined methods<br>• Other | • Researchers<br>• Students<br>• Mentor/RCR Instructor<br>• Institution<br>• Industry<br>• Editors<br>• Participants/Public/Media<br>• Peer-reviewers<br>• Policy makers<br>• REC/IRB<br>• Clinical professional<br>• Research integrity officer<br>• General | • Survey or assessment<br>• Interviews<br>• Focus group/expert panel<br>• Published material<br>• Retractions, notes, or bibliometric data<br>• Industry document<br>• Guidelines, policies, or university requirements<br>• Teaching material<br>• CV and application forms<br>• Case reports<br>• Combined sources<br>• Other | • Determinants<br>- System<br>- Personal<br>- Awareness and compliance<br>• Problem/State of affair<br>• Approaches<br>- System<br>- Awareness and compliance<br>• Consequences<br>• Research on RI | • Describe, explore, or quantify<br>• Test a hypothesis<br>• Denounce or detect QRP and misconduct<br>• Build approach<br>• Assess approach efficacy<br>• Build or validate research tool |
|  | • Not Empirical<br>• Systematic reviews and meta analyses | — | — | — | — | — |
| To identify the *topic of interest*, we looked more particularly at the title and abstract to highlight what the authors appeared most interested in discussing. There could be overlap or different interpretations in this category, but it provides a general overview of the themes most addressed in the field. | We classified included papers according to whether or not they reported empirical work, as defined earlier. A few systematic reviews were also found but we did not include them in further classification processes. | This category shows the general methodology used in the investigated paper. Naturally, it will be tightly linked with the *source of data* | Here we show who participated or was studied in the work. Depending on the *source of data*, these actors might not have been investigated directly, but by a proxy (e.g., Looking at researchers' misconduct through retractions, looking at editors' policies through guidelines for authors, etc.) | In the source of data, we showcase the type of data that was gathered to perform the research by showing from which source it was acquired. | Here, we identified which particular step of the integrity problem is studied (i.e., the determinants or causes for misconduct, the problem itself, approaches to fight the problem, consequences of the problem, or research on research integrity). | For each focus, we described what we considered appeared to be the general objective of researchers. |

Except for the ‘determinants’ and the ‘approaches’ subcategories which were weighed, each article was fitted only once in each category. In case of ambiguity, we revised the papers further to determine what the authors highlighted most in the title and abstract, and we decided the classification based on their emphasis. For example, if a paper looked at guidelines and policies for plagiarism, the paper would obviously be a good for for both the Topic class of ‘guidelines and policy’, and the one of ‘plagiarism’. In such a case, we decided according to the terminology used by the authors in the abstract and title. We did not assess the quality of included literature.

We first classified all relevant papers according to their *topic of interest*. We then used the abstract and full text to determine whether the article was empirical or not. For the purpose of our analysis, we considered anything that includes a minimal description of data collection and analysis, from qualitative research to bibliometric studies or textual analyses, as ‘empirical’.

We further classified each empirical article according to (i) the general methodology; (ii) the studied population; (iii) the source of data collection; (iv) the focus of interest; and (iv) the research objective.

In addition, for papers in which the focus of interest was ‘determinants’ of misconduct and QRP, we extracted the specific determinants found in the empirical work and classified them between *personal* issues, issues with the *system*, and issues related to *researcher’s awareness or compliance*. Likewise, in papers in which the *focus of interest* was ‘approaches’ to misconduct and QRP, we classified the approach as either targeting the *system*, or targeting *researchers’ awareness or compliance* (note: we luckily did not find any approaches that proposed to change personal characteristics such as gender or personality, so we did not include *personal* approaches).

After completing the classification, we analyzed our data in Excel to observe ongoing trends. We used the data visualization program Tableau Software inc. to build figures that illustrate our findings.

### Data availability

The full dataset, with both included and excluded records and full classification categories is available in ‘Online Resource 3 - General data’, and the data on determinants and approaches are available in ‘Online Resource 4 – Determinants and approaches’. ‘Online Resource 2 – Instructions for use’ Describes these datasets in greater depth to allow re-use and extension. We welcome future analyses and queries on our data.

## Study limitations and other considerations

Given the current lack of a comprehensive review in the field of RI, we consider our work to be a first step to expose what has been done and how it has been done in research on RI. That being said, as in any research project, several limitations were inevitable to allow us to manage the amount of data gathered with the resources at hand.

First, we decided on a cut off of 2005–2015 to grasp the bulk of research on RI that happened after the impactful Nature paper *Scientists behaving badly* (Martinson et al. 2005), widely recognized as a milestone in the field. As this review was the first step of a bigger project, we had to set a cut off to achieve a realistic record sample. Starting this extractions in 2016, we chose not to include literature published after 2015 since it might not be fully archived on databases at the time where we performed the search. We invite follow ups on our study as it would be very interesting to see what has happened in the most recent years.

Furthermore, we limited our search to records classified as ‘articles’ to obtain a more manageable and relevant subset of research to include in our analyses. Although we are aware that this automatic classification is not flawless (i.e., it sometimes includes editorials, news pieces, etc., and it might overlook a few research articles), we considered this automatic classification to be the best way to obtain a manageable sample of papers in which the bulk of empirical research on RI should be present despite being aware of the costs of this choice. During the manual screening of the papers, we further excluded papers that were evidently not ‘articles’ (e.g., labeled editorials, labeled news reports, short conference abstracts, and letters to the editor). Nonetheless, to avoid biasing our inclusions to the terminology used by journals to distinguish article categories, and because we noticed that empirical data were sometimes reported in differently labeled records, we kept other papers with a more substantial format (e.g., opinions, commentary, viewpoint, ethics corner, correspondence, etc.) when they were automatically classified under the ‘article’ category.

In light of the two former points, and added to the fact that we lacked a reference point, it was difficult to evaluate the completeness of our sample and the sensitivity of our search strategy. Our findings should thus not be considered in isolation of the methods we have used (e.g., search terms included and not included, the way we defined integrity for the purpose of this research, etc.) and choices we have made (e.g., document type, years included, etc.) to reach a manageable sample of papers.

It is also essential to note that a certain level of subjectivity cannot be fully excluded from the classification of the included papers. For example, when looking at the topic of interest, many papers could fit in several topics—a paper on ghost authorship with the industry would inevitably fit into ‘authorship’, but also concern ‘conflicts of interest’, ‘QRP and misconduct’, ‘reporting and publishing’, and so forth. We were careful to select the categories and classifications we considered most appropriate based on what was highlighted by the authors in the abstract. Although the classification process was not triangulated by individual reviewers, uncertainties were marked and discussed until a common agreement could be reached. Oftentimes, we reflected upon, corrected, and revisited our categories to strengthen the fit, but we did so without consideration of trends or hypotheses.

Classifications in one category were also often linked to classifications in another category. For example, papers on ‘RCR training and mentoring’ will often involve ‘approaches’ to deter misconduct, have the objective to ‘assess’ a method, and study researchers, students, or RCR educators. We tried to remain as neutral as possible when classifying our articles by building our classification from the content of the paper rather than from expected trends. Nonetheless, we believe that our results should not be considered in isolation but as a whole in which each category may intertwine with another.

Finally, because we decided not to assess the quality of included papers, we included a wide range of journals and paper standards. Within our inclusions, ten articles were published in journals present on Beall’s list of predatory publishers (note however that five records come from the publisher Frontiers, whose status as predatory publisher is now mostly refuted). Given that Beall estimates that predatory publications accounts for 5-10% of all open access articles (Butler 2013), ten papers in 986 is a small proportion. Nonetheless, given the topic of our review, the fact that not all included articles were open access, and the fact that we conducted our search using databases which already screen for journal quality, we considered that one percent was still worthy of mention.

## Results and Discussion

### Inclusions and exclusions

After screening titles and abstracts for relevance to the topic, we included 986 articles. Table 2 highlights the number of inclusions and manual exclusions (i.e., manually excluded after the initial Excel sheet has been compiled). The complete dataset, with both included and excluded papers and full classification categories, is available in the ‘Online Resource 3 – General Data’.

**Table 2.** Number of included and excluded records.

|  | INCLUSIONS | EXCLUSIONS | REASON FOR EXCLUSION |
| --- | --- | --- | --- |
| **Total** | **986** | **2454** |  |
| Empirical | 342 |  |  |
| Non Empirical | 621 |  |  |
| Systematic review/meta analysis | 3 |  |  |
|  |  | 183 | Accessibility and language |
|  |  | 295 | Article format or year |
|  |  | 190 | Cheating (exclusively) |
|  |  | 426 | Duplicate/incomplete/retracted |
|  |  | 81 | Methods and tools |
|  |  | 406 | Research ethics, bioethics, clinical ethics,<br>or ethical role of universities in society |
|  |  | 873 | Scope irrelevant |

Within the 5453 publications yielded by the initial search (i.e., before peer review) in SCOPUS, 2477 records (44.4%) were classified as ‘articles’. This low percentage of articles compared to other publication formats is atypical for scientific fields (e.g., for medicine and health sciences the proportion of journal article surpasses 90%) but similar to fields of politics and policy (as depicted in the SCOPUS Content Coverage Guide of 2016), which may be more aligned with the type of documents published on Research Integrity.

### Empirical coverage: a small proportion of empirical work

Around a third of the included publications described empirical work (N=342; see Table 2). This is surprising given that we only included publications automatically classified as ‘articles’ and thus excluded most editorials, letters, and other more theoretical types of publications. Within our inclusions, theoretical approaches, narrative reviews, recommendations, and opinions were most common (classified as non-empirical in the ‘Online Resource 3 – General Data’).

### Topics of interest: general topics, integrity teaching, and publication ethics are most often studied

We extracted the topics of interest of all included papers and grouped them in categories. When papers were not clearly targeting a specific topic, we classified them in the more general categories of ‘QRP and misconduct’ or ‘Research integrity’, accordingly. Most papers targeted ‘QRP and misconduct’, but a substantial proportion of papers also targeted ‘RCR training, education, and mentoring’; ‘Publication ethics’; and ‘Conflicts of interests’ (Fig 2). The proportion of empirical articles was highest for topics of ‘Cheating and academic misconduct’ (77%), ‘CV and application misrepresentation’ (79%), ‘Research on research integrity’ (57%), and ‘RCR training, education, and mentoring’ (49%). This might result from the relative ease to build empirical designs in such topics compared to others.

**Figure 2.**
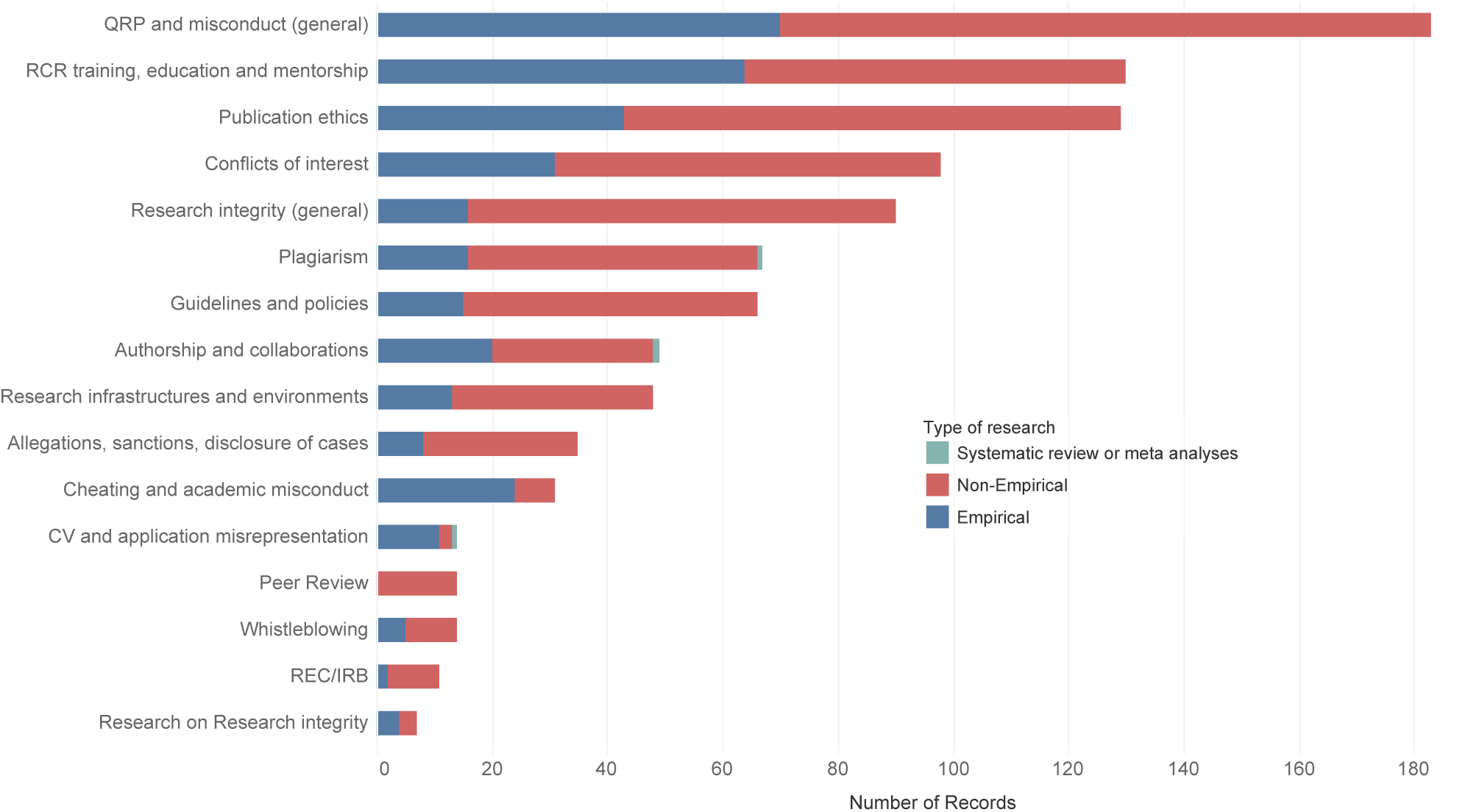
Topics of interest of included papers and corresponding research type. Most papers targeted ‘Questionable research practices (QRP) and misconduct’, but a substantial proportion of papers also targeted, ‘Publication ethics’, ‘RCR training, education, and mentoring’, and ‘Conflicts of interests’. The proportion of empirical articles was higher certain topics, for example ‘Cheating and academic misconduct’ and on ‘RCR training, education, and mentoring’

Although we admit that our keywords may have played a role in the topics found in our results, certain topics were seldom explored in our sample despite their direct relevance towards RI. For example, Research ethics committees/Institutional review boards (‘REC/IRB’), ‘Peer review’, and ‘Whistleblowing’—which may be considered as potential safeguards of integrity in research—were very rarely the main topics of included papers. The current focus, instead, appears to be motivated by describing the problem (‘QRP and misconduct’), strengthening reporting standards (‘Publication ethics’, ‘Conflicts of interest’, ‘Plagiarism’, and ‘Authorship’), and examining integrity training and policies (‘RCR training, education and mentorship’ and ‘Guidelines and policies’).

### Methodologies: surveys and interviews prevail

Over half of empirical papers used direct approaches, such as surveys, questionnaires, interviews, and focus groups (N=175) to obtain their data. Bibliometric studies (N=58) and content and textual analyses (e.g., policy documents, case studies; N=50) were also frequent. These findings corroborate past findings from literature on academic integrity (Macfarlane et al. 2014). The distribution of methodologies alongside more specific research objectives can be seen in Table 3. The more precise sources of data used for each record are available in the full dataset in the ‘Online Resource 3 – General Data’.

**Table 3.**
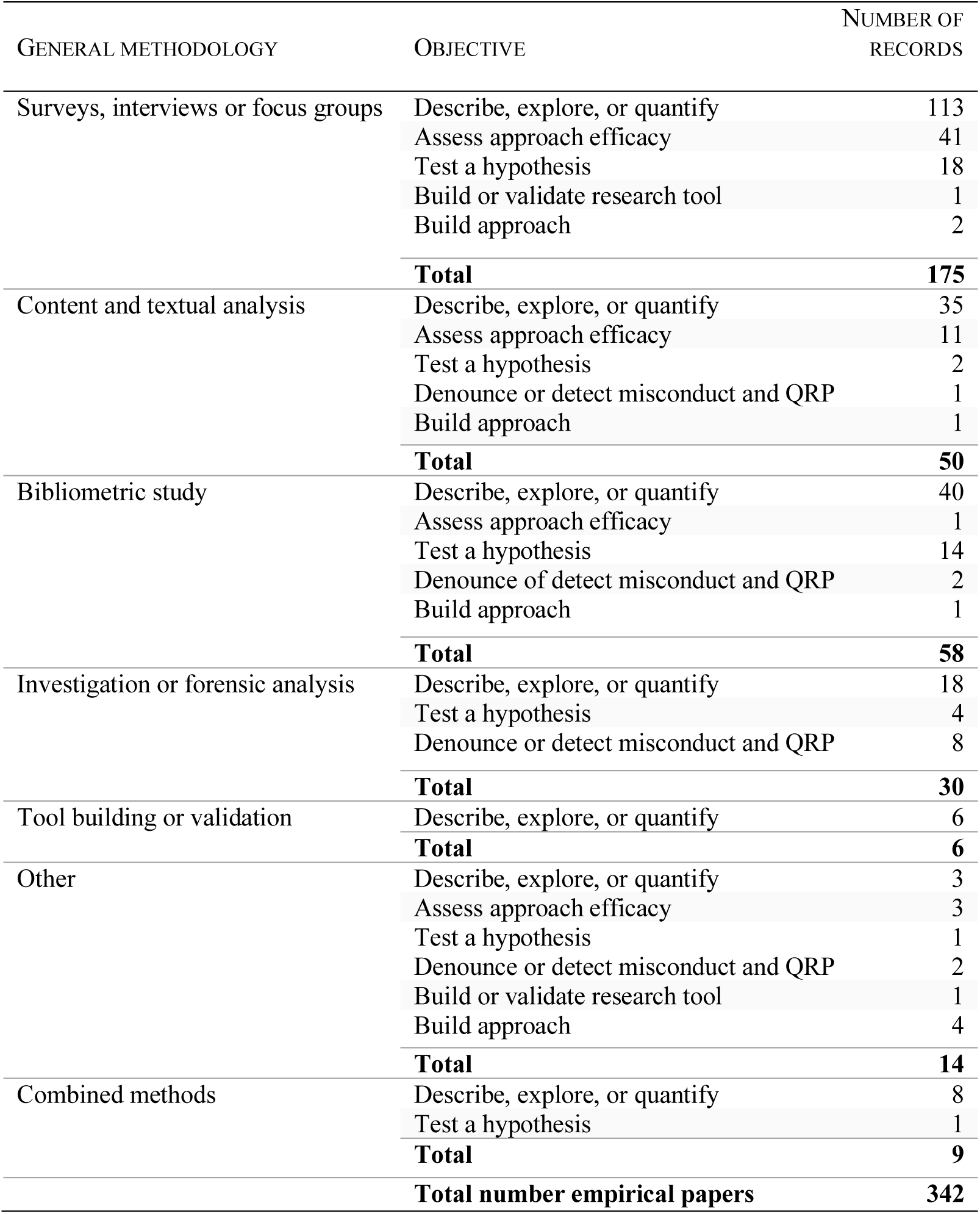
Distribution of methodologies alongside more specific research objectives of empirical papers. About half of all included empirical papers used direct approaches to describe or quantify issues related to research integrity. A fair proportion also used content and textual analyses and bibliometric studies, also mostly to describe or quantify integrity issues

### Studied population: Researchers are most studied, other actors are overlooked

Over 60% of empirical papers study researchers and students, while fewer articles involved actors other than researchers (Fig 3). Researchers and students further account for over 75% of articles that used *direct approaches*—approaches in which investigators directly addressed the studied population, such as interviews, survey, focus groups, and direct observation. Other research actors were most often studied by proxy through documents, reports, or published material.

**Figure 3.**
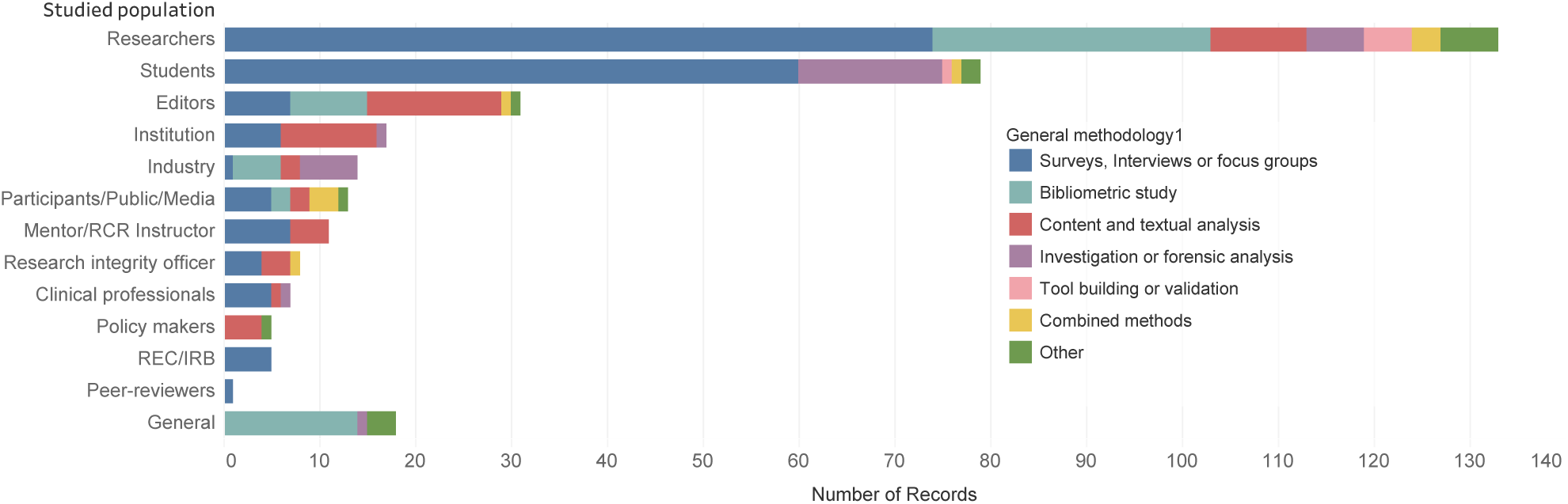
Studied population and general methodologies used. Representation of the major actors studied in included empirical papers. The different colours represent the general methodology used. Over 60% of included empirical papers studied researchers and students—a percentage which rose to 75% within direct approaches such as surveys, interviews, and focus groups (blue bar)

The high representation of researchers and research students is not surprising given that researchers are amongst the most directly affected by and targeted in research misconduct and questionable research practice. Nonetheless, others players involved in research who also have an important role in promoting integrity appear left out from empirical work on RI. Studies on *policy makers* and *institutions*, for example, were sparse and rarely involved direct contact with these actors, despite their crucial role in defining funding and regulations. *Research ethics committees* and *peer-reviewers* were also rarely studied, despite their potentially powerful role in preventing and detecting misconduct. The *public and research participants* were only studied in a few papers that explored the consequences of misconduct (e.g., loss of public trust, risks to research participants), and they were rarely approached directly.

Altogether, this imbalance points to an important gap of knowledge in research on RI. Different members of the research community are unlikely to have the same perceptions and expectations towards research (Bird 2010). Involving a more balanced share of non-researcher research actors would likely bring new perspectives to the discussion. But beyond individual actors’ perspectives, the social contexts and the interaction between actors was also largely untouched by empirical works. Given the complex relationships between research actors and their interrelated dependencies, considering the broader social contexts, the conflicting perspectives, and the shared expectations of different research actors may be essential in building a realistic and comprehensive understanding of RI and misconduct.

### Focus: the emphasis is on the problem rather than on its causes, approaches, and consequences

To map the most studied aspects of RI, we classified all empirical papers according to their focus on the RI issue (i.e., which particular step of the integrity problem they looked at). We defined five general focuses, namely, (i) the ‘determinants’ of misconduct, (ii) the ‘problem/state of affair’ of issues of research integrity, (iii) the ‘approaches’ meant to deter misconduct and promote integrity, (iv) the ‘consequences’ of misconduct and QRP, and (iv) tools and approaches specific to ‘research on RI’. We then further classified the specific research objective we could grasp from the methodology of the paper (see Table 1).

Fig 4 shows that over 45% of empirical work on RI focused on the problem, generally with the objective to describe, quantify, or explore the issue. About a third of the articles focused on approaches to promote RI or deter misconduct, with over half of those assessing the efficacy of an approach. Only 13% of the papers focused on determinants of research misconduct and QRP, generally attempting to test the relationship between a hypothesized determinant and reported misconduct or QRP. Finally, very few articles focused on the consequences of misconduct and QRP (e.g., loss of public trust, risks to research participants, financial waste), or on research on RI.

**Figure 4.**
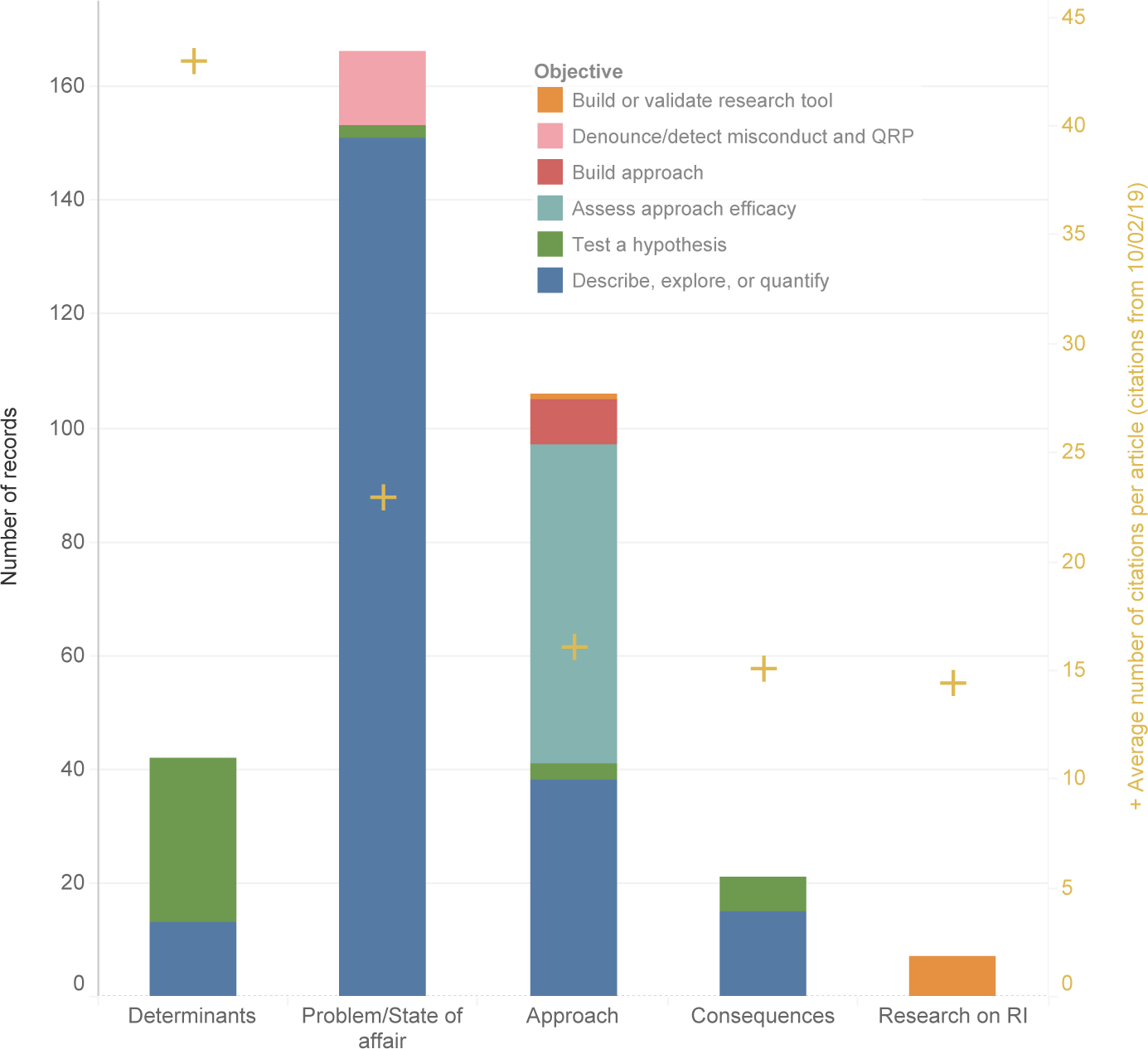
Focus of included empirical work. We classified papers according to their focus (i.e., which particular step of the integrity problem they looked at), and associated research objectives. Yellow crosses show the average number of citation per article for each different focus. We can see that almost half of all empirical work targets the problem while very little articles focus on determinants and consequences. Nonetheless, determinants yielded higher average number of citations (red crosses) than other focus

At first glance, our results insinuate that we know a lot about the *problem* of integrity, but that our understanding of *why* misconduct happens (determinants), what it *engenders* (consequences), and *what can be done* to promote integrity (approaches) is still limited.

### Determinants and Approaches: a mismatch between what is known and what is proposed

We looked in greater depth into papers focusing on *determinants* and *approaches* for misconduct and QRP.

#### Determinants

We extracted findings from the papers focusing on determinants of misconduct and QRP (40 out of the 42 papers on determinants) and grouped them into factor categories to highlight what they found as potential causes for misconduct and QRP.

We then grouped these into broader groups as either highlighting (i) *personal* issues, (ii) issues with the *system*, or (iii) issues with researchers’ *awareness and compliance*. In addition, we computed a weighed indicator for the determinant groups to ensure that regardless of the number of determinants found per paper, each article would account only for ‘one’ paper. (e.g., if a paper found three determinants, each determinant would have a weight of .33 in the paper count). As shown in Figure 5, 45% of the papers found that problems with the *system* played a role in misconduct and QRP, while only 16% of papers found that problems of *awareness and compliance* of researchers were at play.

**Figure 5.**
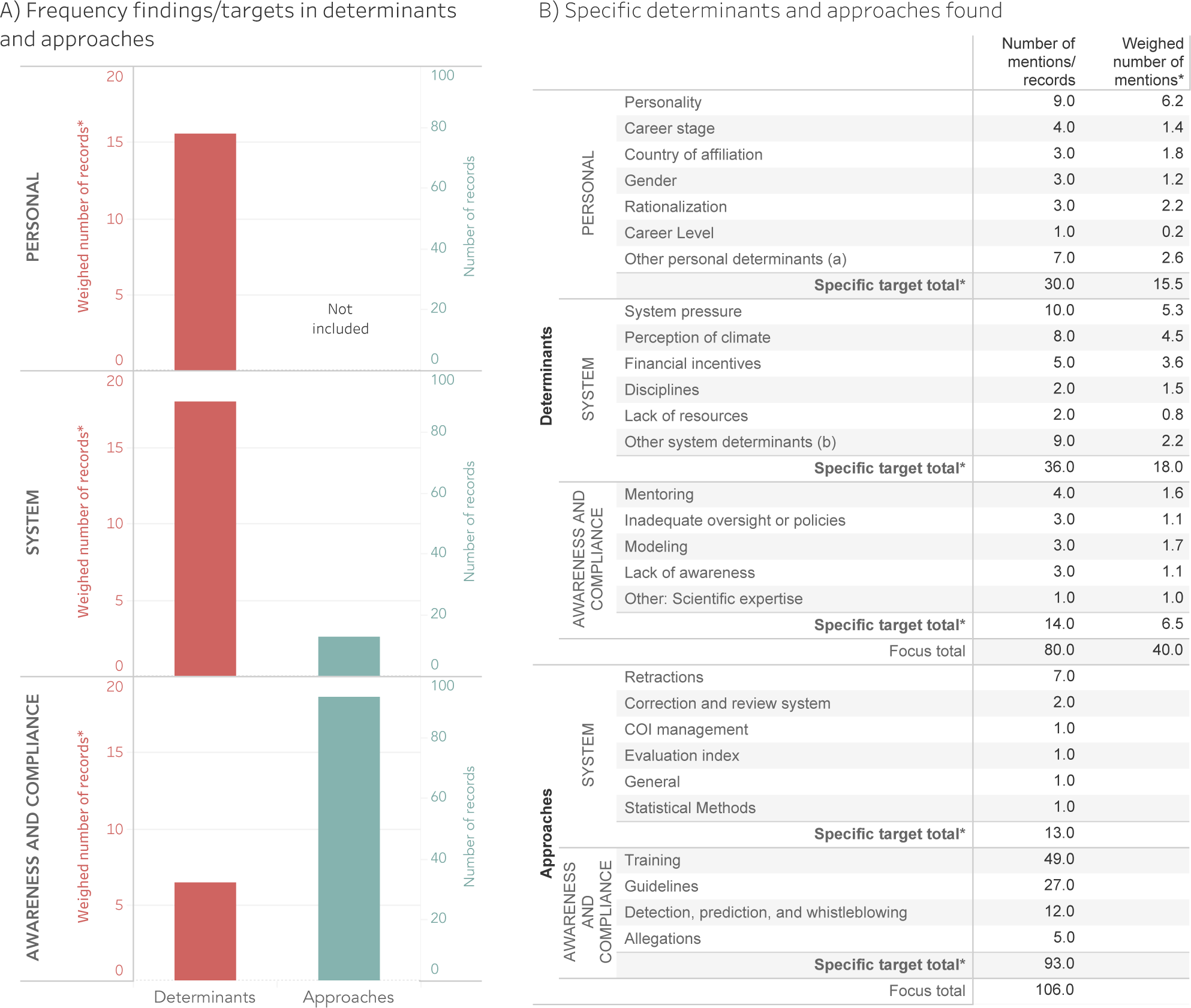
Determinants and approaches to misconduct and Questionable Research Practices (QRP) Determinants (in red; N=79 determinants in 40 papers), and approaches (in blue; N=106 approaches in 106 papers) to misconduct and QRP found or proposed in empirical papers. We can see that most papers on determinants found that issues with the system contributed to misconduct and QRP, while most articles proposing or assessing approaches target researchers’ awareness and compliance Notes: * We equally weighed the determinants to ensure that, regardless of the number of determinants found, each article would account only for ‘one’ paper (e.g., if a paper found three determinants, each determinant would have a weight of .33 in the paper count) † Other personal determinants: Need for recognition, Opportunistic (Internet), Prior misconduct, Single authorship, Personal problems ‡ Other system determinants: Professional relationships, Fear of retaliation, Culture of compliance, Hampered criticism, Type of institution, Job insecurity

Nevertheless, two precisions are important here. First, the papers we classified in the ‘determinants’ categories sometimes reported direct effects on the prevalence of misconduct and QRP, but other times they reported the influence, or the perceived influence of different factors on ethical behaviours, compliance, or reporting bias. Second, even though Figure 5 only includes factors which were found to influence integrity, a few negative or integrity-promoting findings were also highlighted within the papers that looked at ‘determinants’, and some of those effects are not visible in Figure 5.^1^

#### Approaches

Similar to the determinants, we classified papers which targeted approaches to misconduct and QRP into categories which we later grouped as either targeting the *system*, or targeting researchers’ *awareness and compliance*. We did not include *personal* issues in the approaches, as we considered these to be somewhat immutable (i.e., no approach can really target or aim to change gender, seniority, discipline, or country of affiliation). As we can see in Figure 5, almost 88% of papers on *approaches* targeted researcher’s *awareness and compliance*, while very few papers targeted the *system*.

The lack of research on determinants of misconduct and systemic approaches for promoting integrity is not new and has been called before (see for example Fanelli 2015), but our findings highlight that this imbalance also reveals a mismatch between what we know may predispose to inadequate research practices and the approaches to target misconduct that are discussed in the empirical literature. Specifically, factors identified as contributing to misconduct and QRP (i.e., determinants) most often point to the *system*, while approaches to deter misconduct and QRP most often target researchers’ *awareness and compliance*.

Additionally, although a substantial number of articles assess approaches to promote awareness and compliance towards responsible conduct of research training, Ana Marušić and colleagues (2016), who performed a Cochrane review looking at the effectiveness of interventions to prevent misconduct and promote integrity in research concluded that “Due to the very low quality of evidence, the effects of training in responsible conduct of research on reducing research misconduct are uncertain” (p. 2).

In sum, although the empirical literature of research on RI provides substantial information on the problem and state of affairs of misconduct and QRP, less is known about potential causes, approaches, or consequences. It is important to mention, however, that many non-empirical articles proposed approaches — also often systemic approaches — to debunk misconduct and QRP or discussed its potential causes. Our findings, however, indicate that even though opinions on causes and ideas for change abound in the field, these ideas often remain untested empirically.

### Geographic distribution: North America is the biggest player, but Europe and Australia are catching up with impact

Affiliations from the United States largely accounted for over half of the literature on RI captured by our sample (Figure 6B). The UK, Australia, Canada, India, Croatia, and the Netherlands followed respectively, each accounting for more than ten included articles. The predominance of the United States is not surprising given that they are the biggest player in published literature worldwide (see for example Phillips 2016). Nonetheless, China, which is rapidly becoming the second most important player in scientific publication, only accounts for seven articles included (0.7%). It is possible that the language limitations of our study contributed to this disparity as we only included English language articles, yet it is surprising that such upcoming players generate so few research on RI aimed to be captured by and influence the international discourse.

**Figure 6.**
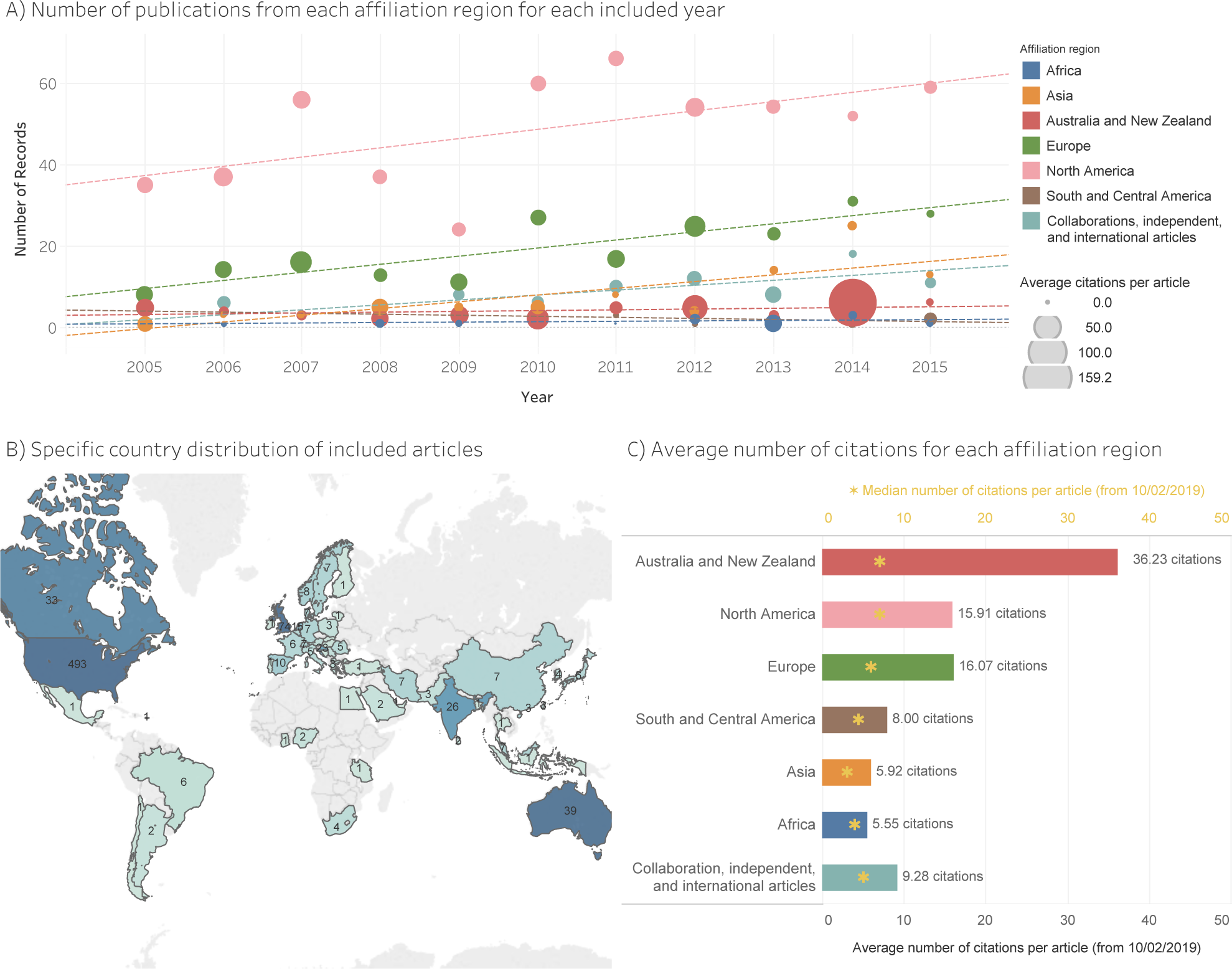
Citation and publication distribution. (A) Number of article included per affiliation region for each included year. The size of markers represents the average number of citations per paper acquired by such region in a specific year. Trend lines illustrate the overall Research on RI publication growth for each region. (B) Specific country distribution of included articles. We did not include collaborations (i.e., articles with several countries mentioned in the affiliations included in the reference record), international (i.e., articles in which the main author was represented by an international organisation), and independent articles (i.e., where the main author did not mention a located affiliation) on the map. (C) Average number of citation per paper for each affiliation region. Australia/New Zealand dominates the average number of yearly citations per article, but this is due in part to one heavily cited paper. The yellow asterisks, which display the median number of citations per paper, show much greater uniformity between continents.

### Citations analysis

Although we did not do a thorough citation analysis, we wanted to have an overview of the citation patterns of included articles. A few things are important to note before getting into our findings. First, we extracted citation counts directly from the databases (i.e., each database counts the citations of its articles based on its pool of included material, and therefore differs), a database effect is thus possible. Additionally, when the citations were not available in the database (i.e., PubMed), we looked for citations by searching for the DOI or title first in SCOPUS, then in Web of Science, and if unavailable, we grabbed the citation counts from Google Scholar. We marked the source of the citation count in the column ‘Citation Source’ in the ‘Online Resource 3 – General Data’. Second, we looked at the total number of citations for each included paper without normalizing for the ‘age’ of the paper. We made this decision to avoid possible issues linked with normalization (Ioannidis et al. 2016). Consequently, it is important to consider that reported citation means and median may be influenced by the number of years the publications have been online, the output of the years following publication, or, on a country level, the size of the output in early years of research on research integrity; we include a figure showing the average citations per paper for each publication year in Figure 6.

We extracted citation counts from SCOPUS and Web of Science on the 10^th^ of February 2019 (older citation counts from 7^th^ November 2017 are also available in the Online Resource 3 – General Data). On average, articles were cited 15 times, yet the distribution of citations was heavily skewed. The median number of citation was 6, and 103 articles (10%) were seemingly never cited February 2019. Over half of the total citations came from less than a tenth of the included papers (7.6%). Looking specifically within empirical papers, we further noticed that articles focusing on *determinants* of misconduct and QRP yielded on average more citations per paper than research focusing on the problem, its approaches, or its consequences (see the yellow crosses in Fig 4). When looking at highly impactful papers (we selected a cut off of 30 citations; N=77), 64% were empirical, and over half (54.7%) had a main affiliation from the United States.

When looking at the citation weight for different continents, it was clear that North America generates most citations in research on RI, but that this dominance of citations is partly due to the important number of publications North America generates (especially the United States). In fact, North America has a lower citation average than Australia/New Zealand, and a citation average similar to European averages (Fig 6C). Australia/New Zealand has the highest citation average, but this may be due to one very highly cited paper. The median number of citations per papers (i.e., the yellow asterisks in Fig 6C) are more uniform between continents, ranging from a median of 7 citations (i.e., North America and Australia/NewZealand) to one of 3 citations (i.e., Asia).

The skewness of citation distributions is not specific to research on RI and often occurs within single journals (see for example Larivière et al. 2016). Nonetheless, the highly-skewed distribution of citations included in our research proposes that research on RI is not immune to such dynamics.

Within the 10% of the literature that was never cited, we noted that only 25% were published in 2015. It thus seems unlikely that these records are simply cases of ‘Sleeping Beauties’ (van Raan 2004), and the uncited records probably have slim chances of being taken up in the future.

In light of these findings, two questions came to mind: first, are there more efficient dissemination systems that could ensure utility and uptake of research on RI; and second, is it possible that published research on RI is being used but not attributed as such? It is conceivable that a significant part of the readership of research on RI uses RI literature to stay up-to-date, to gain insight, and to update training or policy rather than to conduct research, thereby using the findings without citing the articles as such?

We have not conducted a deeper analysis about the source of the citations and about possible network in the citing patterns, however we assume that such analyses may yield interesting results. In particular, investigating whether citation counts of research on RI correlate with implementation, systemic changes, and policy building may be relevant to better understand dynamics and impact in the field in the future.

## Conclusions

Research on RI is a field that is difficult to review systematically. The lack of consistency of its key terms, the absence of a clear delimitation of its scope, the interdisciplinary nature of the journals it targets, and the inconsistency of the article formats it employs to report empirical works make research on RI a fractionated field in which systematic and comprehensive overviews are challenging.

Nevertheless, our analysis of a decade (2005-2015) of scientific articles in the field of research on RI reveals that the body of literature on RI is growing in size and importance, and that the field is still largely dominated by non-empirical papers. While general topics of misconduct, questionable research practices (QRP), and reporting standards are substantially addressed, other relevant topics are largely overlooked. Researchers and students are most often studied empirically but other research actors, such as funders, policy makers, research administrators, are rarely involved.

Empirical research tends to describe the *problem* (e.g., misconduct prevalence) rather than its potential *determinants, approaches*, or *consequences* The few articles looking at determinants most often link the research *system* to bad research practices, while articles assessing approaches to prevent misconduct and QRP most often targeted *researchers’ awareness and compliance* rather than *system* changes.

Even though the present work is only a first glance in the broad and comprehensive body of research on research integrity, several points brought up by our analysis may serve to inspire future research agendas.

First, the sense of urgency attached to the topic of misconduct appears to push scientists to explore new venues for solutions rather than to optimize pre-existing opportunities. In fact, past research on RI most often discussed problems with research integrity and reporting standards, and responded by proposing new surveillance and compliance techniques. Nonetheless, in focusing on new approaches, past research may overlook important safeguards which are currently available in the research organisation (e.g., peer-review, whistleblowing, research ethics committees). Grasping and optimising these existing opportunities might be a relevant target for future research and interventions on RI.

Second, although the predominant involvement of researchers and research students in research on RI is justified given their implication at the core of research practices, involving participants beyond research-producers in future research on RI might help broaden our understanding of the problem. In particular, exploring the perspectives of different research actors and the social context that links these actors might help assess the possibilities, impact, and acceptability of different approaches to foster integrity. In the same way, involving topics and actors who play important roles early in the research process (e.g., research ethics committees, policy makers, funders) may be key to better understand how misconduct can be prevented.

Finally, reconnecting the approaches that are proposed and assessed empirically to what is already known from past research on determinants of misconduct may be essential to increase the success of future approaches to foster RI and deter misconduct. In other words, research on research integrity may need to assess feasibility and success of systemic approaches beyond researchers’ compliance and awareness.

In sum, research on RI undertaken over the past decades has undeniably produced useful knowledge and improved our understanding of the issues faced by researchers and the research system, and it certainly continues to do so. Our review highlights the areas, methods, and actors that have been most studied, and sheds light on points which have been overlooked. Being aware of unanswered questions in research on RI is a first step toward generating executable knowledge that will allow us to better align the research agenda with the goal of promoting integrity in research.

## Supporting information

Online Resource 1 - Building the classifications

Online Resource 2 - Instructions for Use

Online Resource 3 - General Data

Online Resource 4 - Determinants and approaches

Online Resource 5 - Peer Review Reports

## Acknowledgements

We want to thank Prof. Raymond De Vries for his help in revising the manuscript. We would also like to thank the organisers of the Doctoral Forum of the 5th World Conference of Research Integrity (Nicholas H. Steneck, Elizabeth Heitman, and Nils Holger Axelsen) and its participants for their comments and recommendations regarding this work. We also thank peer-reviewers from PLOS One and from Science and Engineering Ethics for their useful comments which helped improve our manuscript.

## Online Resources

### Online Resource 1 – Building the classifications

Online Resource 1 – Building the classifications gives additional information on the process used to analyse and classify the articles included in our study.

### Online Resource 2 – Instructions for Use

Online Resource 2 – Instructions for Use simply describes how Online Resources 3 and 4 should be used, and clarifies each of their columns.

### Online Resource 3 – General Data

The ‘Online Resource 3 - General Data’ csv file contains the original results extracted from SCOPUS, Web of Science, and PubMed to which we added columns with variables classifications. We preserved essential elements of from the database export to allow future use of the data. We use EID as unique identifiers, and the EID can be used in their respective databases to retrieve each record.

### Online Resource 4 - Determinants and approaches

In the ‘Online Resource 4 - Determinants and approaches’ csv file, we copied all records which focused on *determinants* or on *approaches*. We included those records in a new sheet because some articles on determinants found more than one potential factor to contribute to misconduct, and we needed to separate this information on multiple lines. We use EID as unique identifiers, and the EID can be used in their respective databases to retrieve each record.

### Online Resource 5 – Peer Review Report

## Footnotes

1 When training was found to deter misconduct and QRP, (i.e., to promote integrity) we counted it as if the paper stated that ‘lack of awareness’ contributed to misconduct and QRP (e.g., Kraemer Diaz et al. 2015; Geller et al. 2010). When papers found no effects of potential factors, we did not include those factors in the findings in Figure 5. The negative effects found were as follows: Wolfgang Stroebe and colleagues (2012) found that social psychology (i.e., discipline) was *not* more prone to fraud than other disciplines. Karen Woolley and colleagues (2011) found that, although country of affiliation, prior misconduct, and single authorship were related to higher misconduct-related retractions, declarations of financial incentives were not. Michael Mumford and colleagues (2007) found a whole array of factors encompassing all three categories of *personal, systemic,* or *awareness and compliance*, but they also found that work commitment and limited competition did not promote unethical decisions. Finally, Daniele Fanelli and colleagues (2015) found that inadequate oversight or policies, financial incentives, hampered mutual criticism, and career stage affected scientific integrity, but not gender and pressure.

