## Supplementary material for "A decade of empirical research on research integrity: what have we (not) looked at?": Online Resource 1 - Building the classifications

To build the classifications for our research, we used an inductive process (Elo and Kyngäs) based on the findings from the first set of papers retrieved (i.e., the SCOPUS search). An inductive process means that we started with the general idea of describing research, and that we decided on which categories and classification options we should include based on what we found in the abstracts and papers assessed. In an inductive process, categories are highly mobile in the beginning of the analysis, but as we carry on the analysis, recurrent themes and groups of themes slowly solidify the nodes that we choose to look into. As the analysis advances, the process becomes more and more deductive, with new nodes being created when new information is not covered in the past nodes. At the end of the process, the nodes are looked at all together, and through axial coding and connection of concepts, merged in fewer nodes.

For our particular work, this is how we proceeded:
We started with the broad, general idea of describing the literature. Initially, we had a column for topic, and a column for whether the study was empirical or not, as these two points were interesting to us from the onset. We classified papers in these two columns, and, when the study was empirical, we wrote, in another column, one or two sentences to describe what the study was about. We rapidly realised that a series of aspects described in the papers were recurrent, namely, the topic, the methodology, the population, etc. We then found that the same methodology (e.g., content and textual analysis) could be used with different populations (e.g., researchers vs. editors), or with a different data source (e.g., retractions, notes, or bibliometric data vs. guidelines, policies, or university requirements). We thus added the 'studied population' and the 'source of data' categories, and added classification options as we continued to review the literature. We also noted that, in the column in which we described the studies, we hinted on the 'objective' of the researchers (e.g., assessing an approach, describing, exploring, or quantifying a problem, etc.), so we added this category. Realising a trend in the 'objective' categories, we then decided to group those objectives in the 'focus', to express whether the study was describing the causes, the problem, the approaches to the problem, or the consequences of the problem. This grouping allowed a better overview of the research that had been done.
In this process, NAB first read and classified the papers. After one first 'round' of classification, she met with WP and discussed the (then very numerous) nodes she extracted. Together, NAB and WP classified these nodes into broader nodes in order to have fewer, more encompassing classifications. NAB then looked back at the specific abstracts and papers to make sure that the new broader classifications grasped the specific differences accurately. In this process, she sometimes had to create new classifications, or to adapt the wording of the classifications to allow for ambiguous papers to fit in. NAB and WP met again, and repeated this process, until both were satisfied with the overview and simplicity that the categories allowed while retaining enough details to yield accurate knowledge. This process was key to building a comprehensive categorisation of articles. For instance, NAB initially had over 160 different 'topics of interest'. After repeatedly sitting down with WP to look at these topic while investigating for more details in the abstracts and the papers, they managed to reduce these topics in the final 17 topic classifications.

It is important to note that inductive coding is, by definition, dependent on the reviewers. In other words, different reviewers may look at the same data and build different categories, but once the categories are built, deductive analysis (i.e., placing the articles in the respective categories) should yield similar results regardless of the coders. Unfortunately, since this project was only a first step to a bigger project, only NAB went through the full coding process. Nevertheless, our goal being to build simple groups which could help us make sense of the broad diversity of the literature on research integrity, we believe that this method served our purpose.
