## Supplementary material for "A decade of empirical research on research integrity: what have we (not) looked at?": Online Resource 2 - Instructions for Use

The ‘Online Resource 3 - General data’ comprises the bulk of the data used in our literature analysis, both included and excluded data
The ‘Online Resource 4 - Determinants and approaches’ contains curated data used in the analysis of the determinants and approaches.

**Online Resource 3 – General Data:**
The ‘Online Resource 3 - General Data’ csv file contains the original results extracted from SCOPUS, Web of Science, and PubMed to which we added columns with variables classifications. We preserved essential elements of from the database export to allow future use of the data. We use EID as unique identifiers, and the EID can be used in their respective databases to retrieve each record.

The classification variables we added are:

- Inclusion/exclusion: Included (I) or Excluded (R) papers
- Reasons for exclusion: For each excluded paper, we mention the reason for exclusion. We left this cell blank for included papers.
- Type: This column mentions the type of research. We classified included papers between 'Empirical', 'Non-Empirical', and 'Systematic review or meta analyses'. For excluded papers, we entered 'Unclassified' as we did not classify the type of research of excluded work.
- Topic of interest: The topic of interest column lists the topics mainly covered in all included articles.
- General methodology: In this column, we classified the general methodology of included and empirical papers.
- Studied population: This column highlight who participated or was studied in the work. This variable is classified only for included empirical articles.
- Source of data: Source of data describes where or how the data was acquired.
- Focus: The focus represents the particular step of the integrity problem that is studied (i.e., the determinants, the problem itself, approaches to fight the problem, consequences of the problem, or research on research integrity). This variable is classified only for included empirical articles.
- Objective: Following the focus, we described what we considered the general objective of researchers. This variable is classified only for included empirical articles.
- Cit 0 to 200: This is the number of citations marked in the databases as of 5th November 2017, excluding for all articles with more than 200 citations (N=4). We marked this variable for all included articles.
- Citation 5/11/17: This is the number of citations marked in the database as of 5th November 2017 for all included articles.
- Citation source: This is the database from which the citation count was obtained.
- Affiliation country: This is the country of affiliation of the first and last authors. If the country was different between the first and last author, we noted the paper as a 'collaboration'.
- Affiliation Region: We classified each affiliation country in broader grouping of affiliated regions. These are based on the United Nations Geoscheme. This variable is classified for all included articles.
- Affiliation continent: We also classified each affiliation countries in the respective continents they belonged to. This variable is classified for all included articles.
- Beall's list: This variable simply notes articles that were published by publishers figuring on Beall's list of predatory publishers. For articles published by Frontier, I wrote 'Debated' as the status of the publisher has been highly debated in the past years..

**Online Resource 4 - Determinants and approaches**

In the 'Online Resource 4 - Determinants and approaches' csv file, we copied all records which focused on *determinants* or on *approaches*. We included those records in a new sheet because some articles on determinants found more than one potential factor to contribute to misconduct, and we needed to separate this information on multiple lines. We use EID as unique identifiers, and the EID can be used in their respective databases to retrieve each record.

This sheet includes the following columns:

- Focus: Detailing whether the paper targeted determinants or approaches.
- Detail: describing the specific determinants of misconduct identified
- Target: detailing the general target identified
- Ratio: is a fraction weight in which we equally weighed the determinants to ensure that, regardless of the number of determinants found, each article would account only for 'one' paper (i.e., for each article, we computed 1/x; x=number of determinants found in the article
- Count Appr: Partial repeat of the 'ratio column to allow graphical representation
- Ratio Det: Partial repeat of the 'ratio column to allow graphical representation
- Count Det: This column gives each found determinant a count of '1' to allow the 'Number of mentions' in Fig. 5
- Note: Note for specific records, such as exclusions or negative findings

For any queries, feel free to contact Noémie Aubert Bonn at
