## Supplementary material for "A decade of empirical research on research integrity: what have we (not) looked at?": Online Resource 5 - Peer Review Reports

### <u>Online Resource 5 – Peer Review Report</u>

As mentioned in the Note from the authors at the beginning of this manuscript, the present manuscript has spent more than a year in Peer Review and has been rejected from the journals PLOS ONE and the Journal of Science and Engineering Ethics.

In order to provide our readers with all the tools available to judge our work and understand its strengths and weaknesses, we wished to publish the comments obtained in the journals where this manuscript has been peer reviewed. We are thankful for the editors of PLOS ONE who discussed our request and allowed us to publish the anonymized review. We are also thankful for the editors of the Journal of Science and Engineering Ethics and the integrity team of Springer Nature for thoroughly discussing our request, but unfortunately, given the closed peer review policy at Springer Nature, we were unable to provide information about the peer review from the Journal of Science and Engineering Ethics.

#### 1. Peer Reviews from PLOS ONE

NOTE: Reviewer reports for this manuscript are posted with permission of the individual reviewers and of PLOS ONE, where the manuscript was peer reviewed, but is no longer under editorial consideration.

**<u>Reviewer #1</u>**: The increasing use of computing power to collect and analyse published articles without a serious attempt to understand the content is to my mind a disturbing trend. This article is a good example of this trend.

The authors report their analysis of almost one thousand articles on the topic of research integrity (defined broadly). Their main finding is that most are not empirical, and that the articles do not deal with actors such as funders of research (no explanation as to why it should).

The authors do not provide an explanation as to why they undertook the research in the first place: Are they concerned about the imbalance between conceptual and empirical research in this area? If so, why?

> <u>Response from the authors</u>: *We clarified that this was an exploratory research to better understand methods and themes targeted in research on research integrity (RRI) given that no comprehensive review of RRI has been done before.*

Is there an explanation as to why most of the articles are in the medical and biological sciences?

> <u>Response from the authors</u>: *We did not address this point in our manuscript. Research Integrity is a field that is overwhelmingly represented in the biomedical sciences for a number of historical reasons which other researchers have discussed. We considered this issue to be slightly beyond the scope of our review and our expertise, so we did not add this information in our manuscript.*

The conclusion of the paper sums up the lack of substance:
"In sum, research on RI undertaken over the past decades has undeniably produced useful knowledge and improved our understanding of the issues faced by researchers and the research system. Nonetheless, our review is a reminder that, like any other research field, research on RI has shortcomings. Remaining reflexive and critical towards this body of research and learning from its shortcomings is a first step toward generating

useful knowledge that will allow us to better align the research agenda with the goal of promoting integrity in research."

If the authors believe that we should be reflexive and critical towards research in RI, they should do so themselves, rather than urge others to do it. And when doing this, they should explain how their results fit into their analysis.

> *Response from the authors: In revising our manuscript, we modified the conclusion to put in front what we really think our research may add to the field by stating the following: "Our review highlights the areas, methods, and actors that have been most studied, and sheds light on points which have been overlooked. Being aware of unanswered questions in research on RI is a first step toward generating executable knowledge that will allow us to better align the research agenda with the goal of promoting integrity in research." In addition, we believe that the many details we added to improve the transparency and thoroughness of our results increased the reflexiveness and awareness we, as authors, addressed to our findings. We hope that these changes sufficiently address this comment.*

**Reviewer #2**: The article is clear in scope and reasonably well written. I feel that the language could be refined further, but I leave it to the authors to decide to do so, because the text is very clear either way.

However, I have a few important concerns that I believe the article needs addressing.

The first concerns the quality of the data. When I tried looking for specific authors in the data set, I almost immediately came across duplicate records, and didn't find articles that I assumed would be included. I don't know what the impact of these imperfections might be on the results, but it seems that the data set needs some further checking and hand-cleaning.

> *Response from the authors: Having duplicates would have been a big issue, but after verifying all the records in our data, we could not come across any duplicate. There may have been a misunderstanding in the fact that the data file contains all 3315 records yielded by our searches, among which we included 955 in our analyses. For each record, we marked whether we included or excluded the record, and if excluded, we marked the reason for exclusion. Many records are thus on the file, yet marked as excluded with 'Duplicate record' as a reason for the exclusion. This may explain the confusion, so we made sure that this was clear in our instructions for use.*
>
> *As for not finding articles assumed would be included, this probably depended on the keywords, titles, and especially article type or our search strategy... Many non-empirical publications on research integrity are published as editorials, notes, etc., which we did not capture in our search. Despite this limitation, we are still convinced that limiting our article types to articles was the only realistic manner to obtain a manageable number of records with the highest probable yield of empirical work. We added several notes in the limitations, in the methods, and throughout the paper to make the costs and limits of this decision fully transparent to the reader.*

The fact that some plausible records were missing is quite likely the consequence of a flaw in the search methods, which the authors discuss a little bit in the text but perhaps have not entirely appreciated. The search was conducted with specific keywords, which almost certainly biased the search towards certain topics and certain kinds of articles.

> *Response from the authors: We appreciate this comment and have made this very clear in the text by adding both in the methods and the limitation sections a note*

*about possible missed records and about the choice of keywords and methods.*

For example, it was not very clear to me why the initial search included terms such as "academic integrity", which usually refer to students cheating at school.

> Response from the authors: *We now specify in our methods that we chose the final keywords after a few adaptations and tests of what would provide a broad and yet specific enough overview of works that have been published on RI. We further explain that in non-English speaking countries, it is not infrequent to see the term 'academic integrity' being used for any form of integrity in the academic context. We thus purposively included the expression 'academic misconduct' and 'academic integrity' despite their more direct relationship to student cheating to allow capturing articles which might have used the term differently to refer to research misconduct."*

The authors state that they excluded studies on academic integrity of undergraduate students "with no apparent extension to RI". However, it seems that some of the papers included are still of ambiguous relevance to the topic. E.g. record 2406 in the data set "undergraduate and postgraduate pharmacy students'…." Is apparently about plagiarizing in the exams. I think the authors ought to give greater details of their inclusion/exclusion criteria, and the logic behind it.

> Response from the authors: *We are thankful for the reviewer for raising this point. In our exclusions, we excluded papers on academic integrity or cheating when they were limited to undergraduate students, or when there was no apparent extension to RI. Since this paper includes PhD students and also briefly compares behaviours considered acceptable for students to those considered acceptable for researchers, we decided to include the paper and classify it in the 'Cheating and academic misconduct' category. We refined our explanation on this matter in the text, by stating further "academic integrity or cheating limited to undergraduate students, or with no apparent extension to RI in the discussion and the abstract of the paper". Indeed, we also included other records when the discussion mentioned that plagiarism in students may potentially influence future research behaviours.*

At the same time, seemingly more relevant terms, like "research bias", or "scientific fraud" were not used in the electronic search.

> Response from the authors: *True. We intentionally did not include the term 'research bias' since it yielded a number of issues which are outside the realm of research integrity, misconduct and QRP as we defined those. In our definition, we looked events in which researchers are (at risk of) transgressing established rules of science in a way that would affect the scientific record, rather than more good research practices or biasing dynamics of research. Extending our search with these terms yielded a number of biases from inadequate statistical models, reporting methods, etc. which were beyond our scope and often beyond reach of researchers.*
>
> *We also agree that 'scientific fraud' is an important keyword we initially missed in our search strategy. We duly performed a new search for the terms 'Research fraud' and 'scientific fraud' and added the results (126 raw new records of which we included 31 articles)to our analysis. We thank the reviewers for catching this!*

A more important issue, however, is that the search was also conducted by restricting, only in some databases, to articles or clinical trials/articles. As some of the results of the study suggest, this approach is likely to introduce important inconsistencies in the search, because the classification by type used in these databases is unlikely to be reliable or consistent. By the way the authors discuss the prevalence of empirical research, I get the impression that they don't entirely appreciate how imprecise and inaccurate, across journals, the classification "article" is.

*Response from the authors: This point is important. Unfortunately, we did not have the resources to analyse all results yielded by a search in which the 'type of record' was left unspecified, since this more than doubled our sample. Nonetheless, we fully explain this issue with honest transparency in the manuscript, making sure the reader is aware of the glimpse of research he/she may get from our work, and the portion which may be missed.*

In short, the search terms and strings used are likely have produced an incomplete and biased representation of the literature. Ideally, the study should be integrated by repeating the search on non-articles, and trying to ensure that the data is complete.

*Response from the authors: See above.*

At any rate, the limitations of the search methods used should be a) justified (e.g., there are too many records if non-articles are included, and give numbers) b) discussed and assessed for its likely impact.

*Response from the authors: We now report the reason for our choices (resources at hand, time, scope of the project as a first step to a PhD thesis, etc.), we detail potential biases such choices might have caused, and we provide numbers of our exclusions in the study flow diagram and in the text.*

Finally, although the classifications and graphics presented were elegant and clear, more details ought to be given concerning the classification criteria for all categories presented in Table 1. Each category should be defined, ideally with details about how decisions of inclusion were made, maybe with justification.

*Response from the authors: This another important catch. We added a row with definitions of each classification in Table 1, noting some particularities each categories might have. In addition, we further described i) how the categories were decided upon in the text, ii) how we built and defined the categories (i.e., Online Resource 1) and iii) what are th options and definitions of each category (Online Resource 2).*

Since the only objective of the article is to present statistics about various types of article and classifications within the field of research integrity, then it is crucial to ensure the highest rigor in how methods are presented, limitations are discussed, and the data is checked for validity.

*Response from the authors: Again, this comment related to the comment by Reviewer 1. We fully agree, and decided to increase transparency to a maximum to ensure that the readers know exactly how our search strategy was performed and what it may have missed. Checking data for validity is very challenging at the moment since this project is one of the first of its kind, and we could not find a reference point to compare or validate our findings. We believe that the work still provides interesting information which may be useful for future research on research integrity, and that the Data file which we attached in Online Resource may prove especially useful for or inspire beginners in the field by locating topics, types of research, and specific methodologies.*

**Reviewer #3**: This is a very informative and timely review. To my knowledge there are no scoping reviews on responsible conduct of research and yet there is obligatory training, scandals and discontent regarding the ethos of scientific practice. Please note that I am not a statistician and cannot review statistics in this paper

The methodology is generally sound barring a few exceptions:
1) there could be more topics included such as social responsibility, ethics of knowledge translation and discrimination in science.

> *Response from the authors: We would have loved to include more topics, especially about the ethics of knowledge, about the social and ethical role of universities and academia, and about research biases beyond the researchers themselves. Unfortunately, such broad topics would have yielded too many records to be able to go into each records and extract the topics, methods, and other classifications with the time and resources at hand (i.e., given that this is only the first portion of a PhD project). In the manuscript, we specified that the resources at hand forced us to make decisions, and we explain the costs and limitations of those decisions with much more details.*

2) many empirical studies are published as commentaries in science journals even though they are full research studies. Even Martinson's study "Scientists behaving badly" in Nature is a commentary. How can you account for this practice? I fear that you may have over-excluded.

> *Response from the authors: True. As mentioned above, we increased transparency and ensured that the readers are aware of what they may be missing in our work. We further added "Although we are aware that this automatic classification is not perfect (i.e., it sometimes includes editorials, news pieces, etc., and it might overlook a few research articles), we used this automatic classification to obtain a manageable sample that was most likely to include the bulk of empirical research on RI. [...] Considering that the impactful Nature paper by Martinson et al. (2005) was itself classified as a 'research note' and not an article, it would be interesting for researchers with enough resources to further this analysis without an initial limit to articles."*

3) Also, you mention that all non-empirical work is not a research article. However, many consider theoretical research to be valid and much more detailed than a commentary. Please define terms used.

> *Response from the authors: We were not certain to understand this point, but we believe that the reviewer refers to the fact that we did not include non-articles to include what we qualify as empirical work. We agree that commentaries may be less detailed and valid than other types of non-article research, and that including only research classified as articles may have missed important, valid, and detailed works. As described above for the other reviewers, we now include a thorough explanation of this choice and transparently discuss the costs and miss that this choice may have yielded.*

The results are well explained and clear with and without tables and figures (although figures are quite interesting for overview). However, you seem surprised that there is more research on researchers regarding misconduct; isn't this somewhat justified especially in the early year of research integrity?

> *Response from the authors: You are right that this emphasis on researchers may be quite natural. In the manuscript, we already included the sentence The high representation of researchers and research students is not surprising given that*

*researchers are amongst the most directly affected by and targeted in research misconduct and questionable research practice.", but we rephrased slightly to ensure that our finding does not indicate surprise, but highlights the potential overlook of other research actors.*

P.18 line 312-314 explains well the results.

*Response from the authors: Thank you.*

The conclusion starts with two sentences that are overgeneralized and unnecessary; the first lines 426-430 should be reworked to summarize the main findings.

*Response from the authors: Right, we agree and took off those sentences.*

Minor comments:
p.1, line 12 : You mention " We found that the body of literature on research integrity is growing in importance, but that the field is still largely dominated by non-empirical publications." However, you do not mention why this is problematic.

*Response from the authors: This sentence is not meant to say that there something problematic with this fact. To make sure the reader does not confuse our intention, we changed 'but' to 'and', so that it expresses the fact rather than a possible lack.*

p.1 line 18-19 You mention that "most empirical work proposing approaches to promoted integrity target the researcher's awareness". Is this always the fact from 2005-2015 ? The more recent literature seemed to shift regarding system based approaches.

*Response from the authors: We would absolutely love to gather more recent papers, but this project was simply a first step from a PhD project and including articles from 2015 onward would be impossible given the resources at hand. We added a note to make this clear and invite future researchers to compare our findings with more recent data: "As this review was the first step of a bigger project, we had to set a cut off to achieve a realistic record sample. Starting this extractions in 2016, we chose not to include literature published after 2015 since it might not be fully archived on databases at the time where we performed the search. We invite follow ups on our study as it would be very interesting to see what has happened in the most recent years."*

p.11, lines 185 you mention paper in Beall's list. It the percentage of papers higher than the average in other fields? I am unsure if ten articles is high or low? Please be explicit about the meaning of this finding.

*Response from the authors: We added a note on this as follows: "Within our inclusions, ten articles were published in journals present on Beall's list of predatory publishers (note however that five records come from the publisher Frontiers, whose status as predatory publisher is greatly debated). Given that Beall estimates that predatory publications accounts for 5-10% of all open access articles (Butler 2013), ten papers in 986 is a small proportion. Nonetheless, since not all included articles were open access, and since our search was performed using databases which already screen for quality, we considered it worthy of mention."*

p.12 table 2; the table has odd characters; however, this may be simply a conversion issue.

*Response from the authors: Thank you for noting this. We believe we have addressed this issue.*
